# Reference-free Analysis of scRNA-seq Data Reveals Elevated rRNA and mtRNA Transcription during Neurogenesis in Axolotl

**DOI:** 10.1101/2024.11.18.624113

**Authors:** Md. Rownok Zahan Ratul, Md. Redwanul Karim, Md. Abul Hasan Samee, Atif Rahman

**Affiliations:** Department of Computer Science and Engineering, Bangladesh University of Engineering and Technology, Dhaka, 1205, Bangladesh; Department of Integrative Physiology, Baylor College of Medicine, Houston, 77030, TX, USA

## Abstract

Analysis of single-cell RNA-seq data is typically performed on a gene expression matrix estimated by aligning reads to a reference transcriptome. However, this approach is difficult to apply to organisms with no or incomplete reference transcriptomes. In addition, events deviating from the reference remain undetected. Here we present a reference-free method to analyze single-cell RNA-seq data based on *k*-mers. We assess the performance of our method on a metastatic renal cell carcinoma dataset and find that it is largely able to capture differentially expressed genes. We then analyze a recently generated dataset to study neurogenesis in Axolotl and observe increased levels of transcription of rRNA and mtRNA during neurogenesis as well as a miRNA with previously predicted links to neuronal development. We also detect lncRNAs and intron retention in heart disease-related genes in diseased cardiomyocytes in an analysis of a congenital heart disease dataset.

## Introduction

Single-cell RNA sequencing (scRNA-seq) has revolutionized the study of cellular heterogeneity, enabling detailed analysis at a single-cell resolution. The standard pipeline for scRNA-seq analysis begins with aligning sequenced reads to a reference transcriptome which is followed by expression quantification to produce a gene expression matrix (1; 2). This matrix serves as the foundation for various downstream analyses, including differential gene expression, gene regulatory network inference, and trajectory analysis (3; 4; 5). The alignment process employed may be a traditional alignment method (6) or a pseudo-alignment technique (7) - the latter significantly improves the speed of quantification.

Despite its widespread use, the standard scRNA-seq pipeline has limitations, particularly due to its reliance on reference transcriptomes. Reference transcriptomes or reference genomes are incomplete or non-existent except for humans and a few other model organisms. For non-model organisms, the lack of comprehensive reference transcriptomes makes scRNA-seq data analysis challenging. Moreover, genetic diversity and population-specific variations can create difficulties and introduce biases in reference-based scRNA-seq analysis of model organisms. Even the human references show a Euro-centric bias, predominantly representing European ancestry (8) and a substantial amount of sequences present in other populations are missing in the reference (9; 10). This bias can lead to mis-quantification or disregard of transcripts resulting in incomplete analysis.

Furthermore, the standard pipeline typically quantifies reads aligned to exonic regions, which are the most expressed sequence regions. While this approach works well under normal conditions, it falls short in scenarios involving disease states such as cancer and tumors, that are associated with deviations from the reference. Events like intron retention, atypical splicing, fusion transcripts, and long intergenic non-coding RNA (lncRNA) transcription, which can trigger various diseases (11; 12; 13; 14), remain undetected in the typical pipeline.

These issues highlight the need for reference-free approaches in transcriptome analysis. In the past, to overcome the limitations of reference-based approaches, reference-free methodologies have been developed for various applications such as mutation identification (15), association mapping (10), analysis of metagenomic data (16). Audoux et al. (17) developed a method based on *k*-mer decomposition to capture exon-intron junctions and intronic regions with differential expression in bulk RNA-seq and Sun et al. introduced an approach for cell type classification with scRNA-seq data (18). Recently, Chuang et al. (19) presented a generalized approach for detecting statistically analogous expressions across samples through anchor-target *k*-mer abundance matrices.

We present a reference-free method for analysis of single-cell RNA sequencing data which can be large and noisy (20). The method is implemented in a tool named scKAR (single cell *k*-mer based Analysis without Reference). It first constructs a non-reference *k*-mer abundance matrix. It then identifies *k*-mers associated with specific cell types or conditions based on a dendrogram of types or conditions. Finally, the *k*-mers are assembled to identify sequences up or downregulated in different cell types, and to characterize transcriptional variability across various conditions.

We find that scKAR can detect important transcripts in non-model organisms, where comprehensive reference annotations are missing. Additionally, it captures non-transcriptomic events—such as intron retention, lncRNA transcription, and microRNA (miRNA) expression—that are often overlooked in traditional analyses. First, we validate our method using a metastatic Renal Cell Carcinoma (mRCC) dataset. We then analyze an axolotl (*Ambystoma mexicanum*) neurogenesis dataset previously generated by Lust *et al*. (21) and uncover elevated ribosomal RNA (rRNA) transcription during early stages of neurogenesis, upregulation of mitochondrial RNA (mtRNA) during peak neurogenesis indicative of the high energy requirements, and detect differential expression of a micro RNA (miRNA) *mir6236* which has been reported to be a key regulator in neural development and metabolism. Moreover, in an analysis of congenital heart disease datasets, our tool successfully detected intron retention and lncRNA transcription in essential cardiac genes linked to cardiomyopathy.

## Results

### A reference-free method for scRNA-seq data analysis

We present a reference-free method for scRNA-seq data analysis which is implemented in a tool named scKAR. Figure 1 shows the overview of our methodology. The method takes as input scRNA-seq data in the form of sequenced reads from a set of cells. First, the method performs *k*-mer decomposition on the raw sequences of the samples to generate a *k*-mer abundance matrix. The matrix is then normalized and *k*-mers present in the reference transcriptome can optionally be excluded. Clustering on the *k*-mer abundance matrix is employed to identify cell types within the samples from which a correlation dendrogram is generated. If a reference transcriptome is available, clustering can be performed on the gene expression matrix instead. Next, *k*-mers with similar variance across different conditions are filtered out using Fisher’s F-test. Removal of each edge in the dendrogram creates a bi-partition of cell types of conditions. A differential expression test is employed to identify *k*-mers that are differentially expressed with statistical significance across each pair of conditions. Lastly, the differentially expressed *k*-mers are assembled de-novo to form contigs that are differentially expressed. The assembled contigs can then be utilized for downstream analyses, such as identifying transcriptional events, including intron retention, splice variants, etc. Additionally, sequences from transcripts yet to be annotated in the species may be studied through homology search.

**Figure 1:**
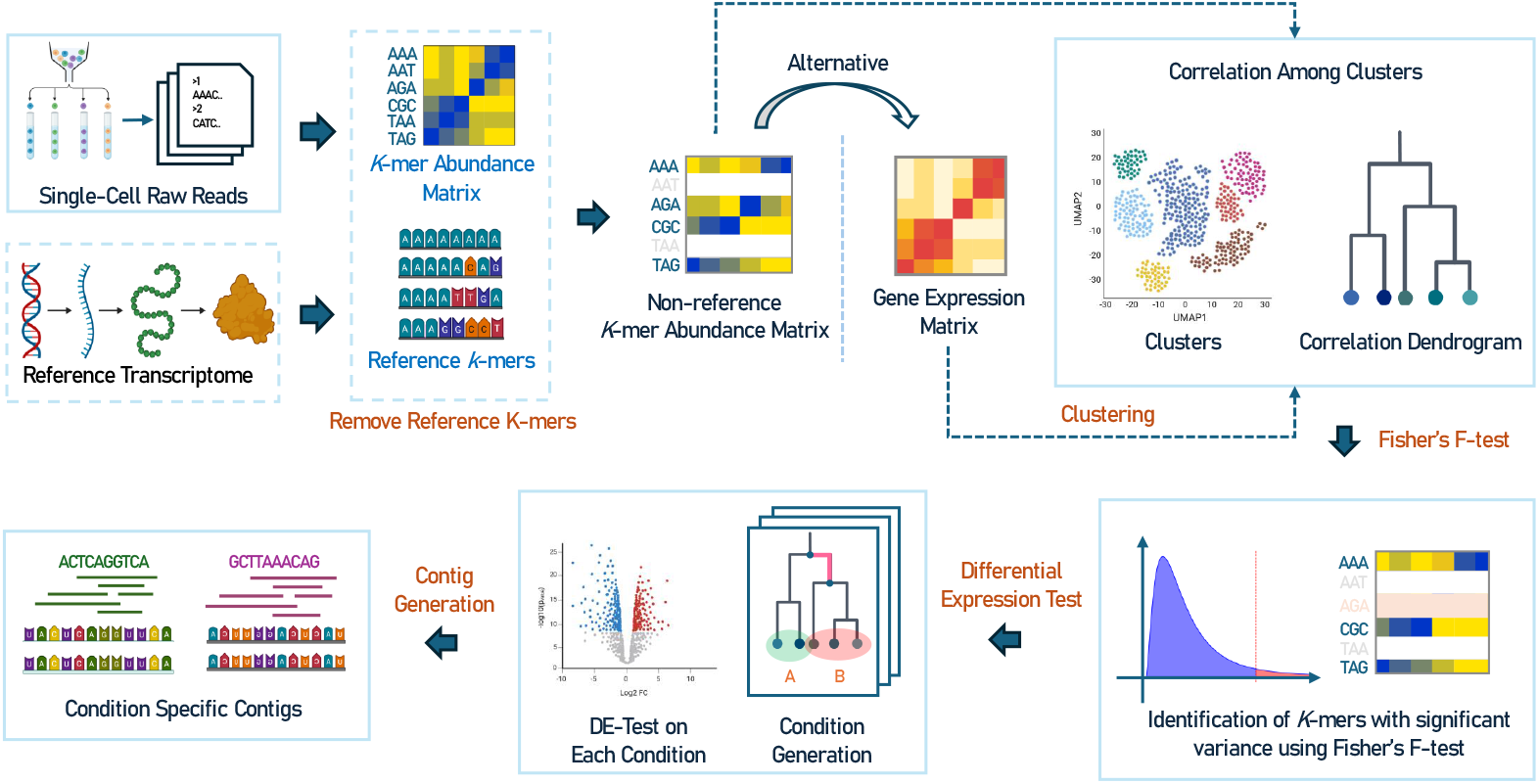
Single cell raw reads are processed to generate a *k*-mer abundance matrix. Optionally *k*-mers presnt in the reference (if available) can be removed from the abundance matrix to yield a non-reference *k*-mer abundance matrix. This abundance matrix (alternatively the gene expression matrix) undergoes graph based clustering to form clusters and correlation dendrograms. Fisher’s F-test is performed on the groups identified from the clustering and the *k*-mer abundance matrix to filter out *k*-mers with similar variance in all the clusters. A differential expression test identifies differentially expressed *k*-mers for each of the partitions generated from the dendrogram. Finally, differenetially expressed *k*-mers associated with different conditions or cell types are merged to generate condition specific contigs.

### scKAR captures differentially expressed genes in a renal cell carcinoma dataset

First, we assess the performance of our method using a Metastatic Renal Cell Carcinoma dataset (22). The dataset comprises single-cell RNA data from 121 samples, including 34 samples of parental metastatic Renal Cell Carcinoma (mRCC), 36 samples of patient-derived xenografts for mRCC, and 46 samples of patient-derived xenografts for papillary Renal Cell Carcinoma (pRCC). Initially, we conducted clustering utilizing the Leiden (23) algorithm on the gene expression matrix of the samples, resulting in the identification of three distinct clusters, as illustrated in Figure 2a. Upon comparing the distribution of samples across clusters with the known sample types, we observed that samples from pRCC were predominantly clustered in cluster 0, while clusters 1 and 2 were mainly composed of samples from parental mRCC and patient-derived xenografts for mRCC. Subsequently, through examination of the dendrogram depicted in Figure 2b, we identified a single bipartition that segregated the cells into two groups. Specifically, one group comprised cluster 0, while the other encompassed clusters 1 and 2. As a result, this bipartition effectively separated the pRCC and mRCC samples into two distinct groups.

**Figure 2:**
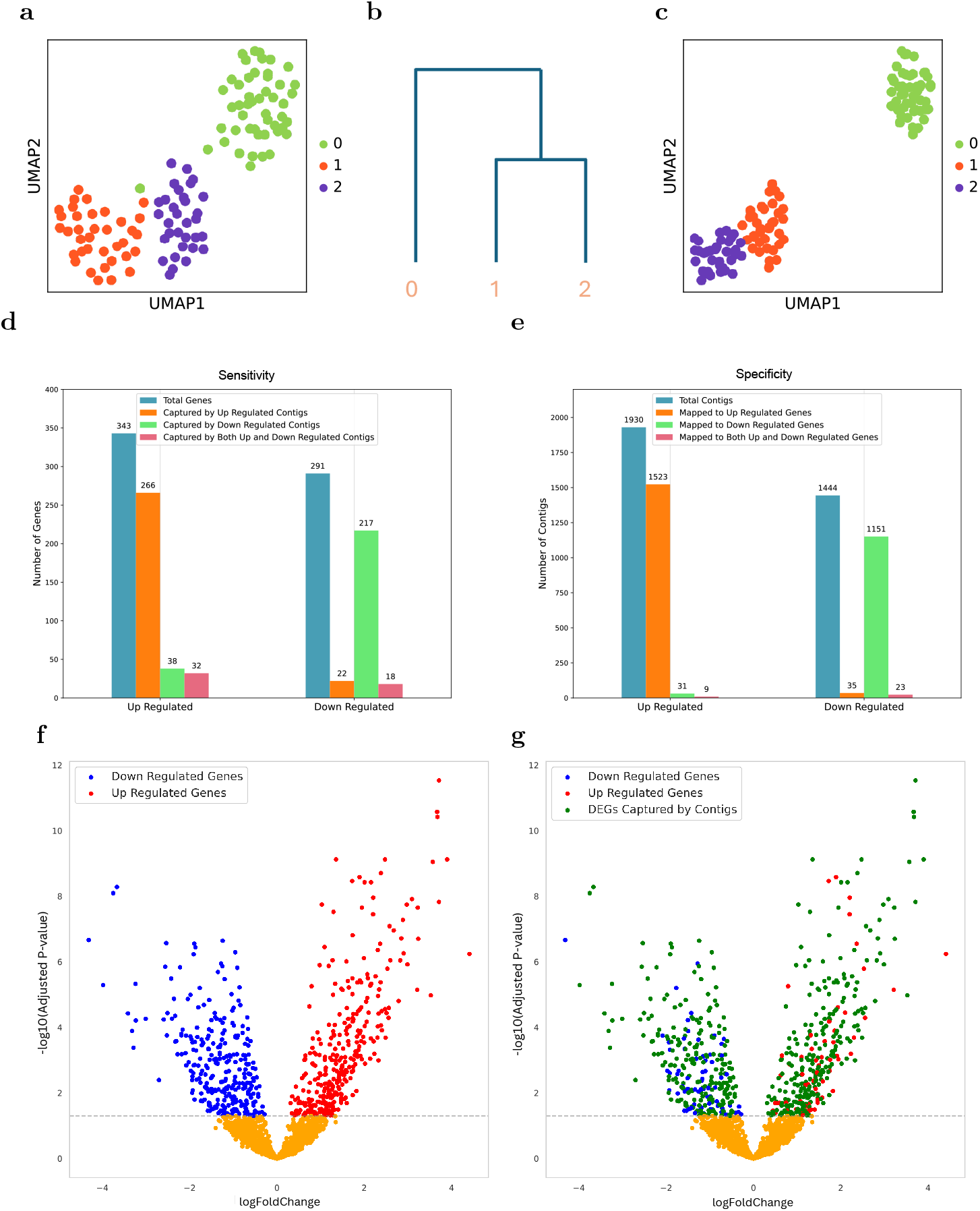
Validation on Metastatic Renal Cell Carcinoma Dataset: **a**. Three distinct clusters corresponding to pRCC, parental mRCC, and PDX-mRCC cells identified by Leiden clustering on the gene expression matrix. **b**. Correlation dendrogram produced from the clustering of gene expression matrix. **c**. Clustering results on the *k*-mer abundance expression domain, achieving a Fowlkes-Mallows index of 0.965 with the clusters on the gene expression matrix. **d**. Sensitivity depicted by bar plots illustrating coverage of DE genes by DE contigs for upregulation and downregulation. **e**. Specificity demonstrated by bar plots showing contigs mapping to differentially expressed genes for upregulation and downregulation. **f**. Volcano plot indicating DEGs meeting an adjusted p-value criterion of 0.05. **g**. Volcano plot outlining the genes covered by contigs generated for validation.

As noted earlier if a gene expression matrix is not available, clustering can be performed on the *k*-mer abundance matrix. We also clustered the count matrix comprising *k*-mers derived from raw reads using the Leiden (23) algorithm, resulting in the identification of three distinct clusters shown in Figure 2c. To evaluate the similarity between the clusters generated from the gene expression matrix and the *k*-mer count matrix, we used the Fowlkes-Mallows index (FM-index) (24). The analysis yielded an FM-index of 0.965, indicating strong agreement between the clustering results. This indicates that *k*-mer abundance matrix clustering is a viable alternative to clustering based on the gene expression matrix.

Subsequently, we employed our method to discern differentially expressed contigs between samples of mRCC and pRCC. As part of our validation process, instead of filtering out *k*-mers in the reference, we applied a filter to exclude *k*-mers that were not present in the reference transcriptome. As a result, we obtained differentially expressed contigs that are present in the reference transcriptome. Following this, we identified genomic regions to which each contig maps using BLAT (25) and ascertained the specific gene associated with each contig mapping employing gffUtils (26) package in Python. Differentially expressed genes were then identified using DESeq2 (27) incorporating the suggested settings from (28) and identifying differential expression with adjusted p-values less than 0.05 where the upregulated genes have log fold change greater than 1 and downregulated ones have less than 1.

To validate our method, we assess both sensitivity and specificity. Here, sensitivity is defined as the proportion of upregulated genes that were captured by upregulated contigs and likewise, the proportion of downregulated genes captured by downregulated contigs. Specificity is defined as the proportion of upregulated contigs that correspond to upregulated genes, and similarly, the proportion of downregulated contigs that correspond to downregulated genes. We compare the differentially expressed genes with the associated genes of differentially expressed contigs to compute sensitivity and specificity. Figures 2d and 2e depict the numbers of DE genes captured by DE contigs and DE contigs mapped to DE genes, respectively. Our method achieved sensitivity of 77.6% and 74.6% for upregulated and downregulated genes, respectively, Besides, specificity of 78.9% and 79.7% was achieved for upregulated and downregulated contigs, respectively. Figure 2f presents the volcano plot of the differentially expressed genes and Figure 2g displays which of these genes are captured by contigs identified as differentially expressed by our method. The sensitivity values indicate that our method successfully detects and captures a large proportion of differentially expressed genes present in the dataset. Conversely, the specificity values show that a substantial majority of the contigs identified as differentially expressed are indeed associated with genes that exhibit differential expression.

### scKAR reveals elevated rRNA and mtRNA transcription during neurogenesis in Axolotl

Axolotl (*Ambystoma mexicanum*), a distinctive species of salamander, is recognized as a model organism for studying regenerative biology (29). Neurogenesis refers to the process by which neurons are generated. This process takes place in the brains of both mammals and axolotls. However, there is a significant difference between the extent, location, and mechanisms of this process between the two groups. Neurogenesis in mammals is most active during the development phase of the brain and reduces with age. Whereas for axolotls, this process persists throughout the postembryonic life. In a recent study, Lust *et al*. generated snRNA-seq and spatial transcriptomics data from axolotl telencephalon to study the neurogenesis process (21). Here, we solely focus on the snRNA-seq data.

The snRNA-seq data includes single nucleus transcriptome obtained from the axolotl brain both under steady-state conditions and after induced injury. snRNA-seq was combined with EdU labeling of S-phase cells to implement Div-seq. Dorso-lateral region of the telencephalon was injured and Div-seq was applied throughout the regeneration process by labeling cells with EdU and collecting EdU^+^ cells at 1, 2, 4, 6, 8, 12 weeks post injury.

We generated two bipartitions: one containing cells in a steady state and the other with cells from the post-injury phase. Using our pipeline, we discovered differentially expressed contigs between these two groups of cells. Approximately 900 contigs were found to be upregulated in steady-state cells, while around 2900 contigs showed upregulation during post-injury. We aligned the contigs to the reference genome of axolotl using hisat2 (30) and identified different transcriptional events as illustrated in Figure 3a. Due to the incomplete nature of the axolotl reference genome, a large portion of the contigs remained unannotated. Specifically, about 75% of the contigs upregulated in steady-state cells and 60% of those upregulated post-injury were unannotated. We performed homology analysis on these unannotated contigs using BLAST (31). Around 8% of the unannotated contigs up regulated during post-injury mapped to ribosomal RNA (rRNA), ribosomal RNA genes, specifically annotated as 12S, 18S, and 28S subunits and mitochondrial RNA (mtRNA) of other species, notably salamanders like *Ambystoma barbouri, Ambystoma opacum, Ambystoma tigrinum* etc, as well as turtles like *Emys orbicularis, Chrysemys picta* and *Malaclemys terrapin*, as mentioned in Supplementary Table S1. On the contrary, less than 1% of the upregulated contigs in steady state aligned to rRNAs. Most of them were predicted rRNA sequences, and very few were experimentally confirmed rRNAs. The alignment of upregulated contigs during post-injury to rRNA suggests upregulation in protein synthesis. rRNA plays an important role in the formation of ribosomes, responsible for translating mRNA into proteins. An increase in rRNA indicates the need for new proteins to rebuild damaged tissue, support neurogenesis, and restore normal cellular functions essential for recovery. On the other hand, the presence of mtRNA highlights the increased energy requirement for cellular activities such as cell division. During regeneration, cells must divide and replace the damaged or lost cells. Upregulation of mtRNAs suggests that the cells are in a high metabolism state, demanding more energy to fuel cell division and tissue regeneration, which are essential for effective healing. Finally, we aligned the contigs to the reference genome of mouse (*Mus musculus*). One of the contigs upregulated during post-injury mapped to the microRNA *mir6236*, which has been shown to enhance neuronal development *in silico* (32), improve insulin sensitivity and reduce metabolic dysfunction (33), and promote skeletal muscle angiogenesis under ischemic conditions (34). These functionalities align with the metabolic and angiogenic demands of axolotl neuroregenesis, highlighting the significance of capturing *mir6236* by scKAR.

**Figure 3:**
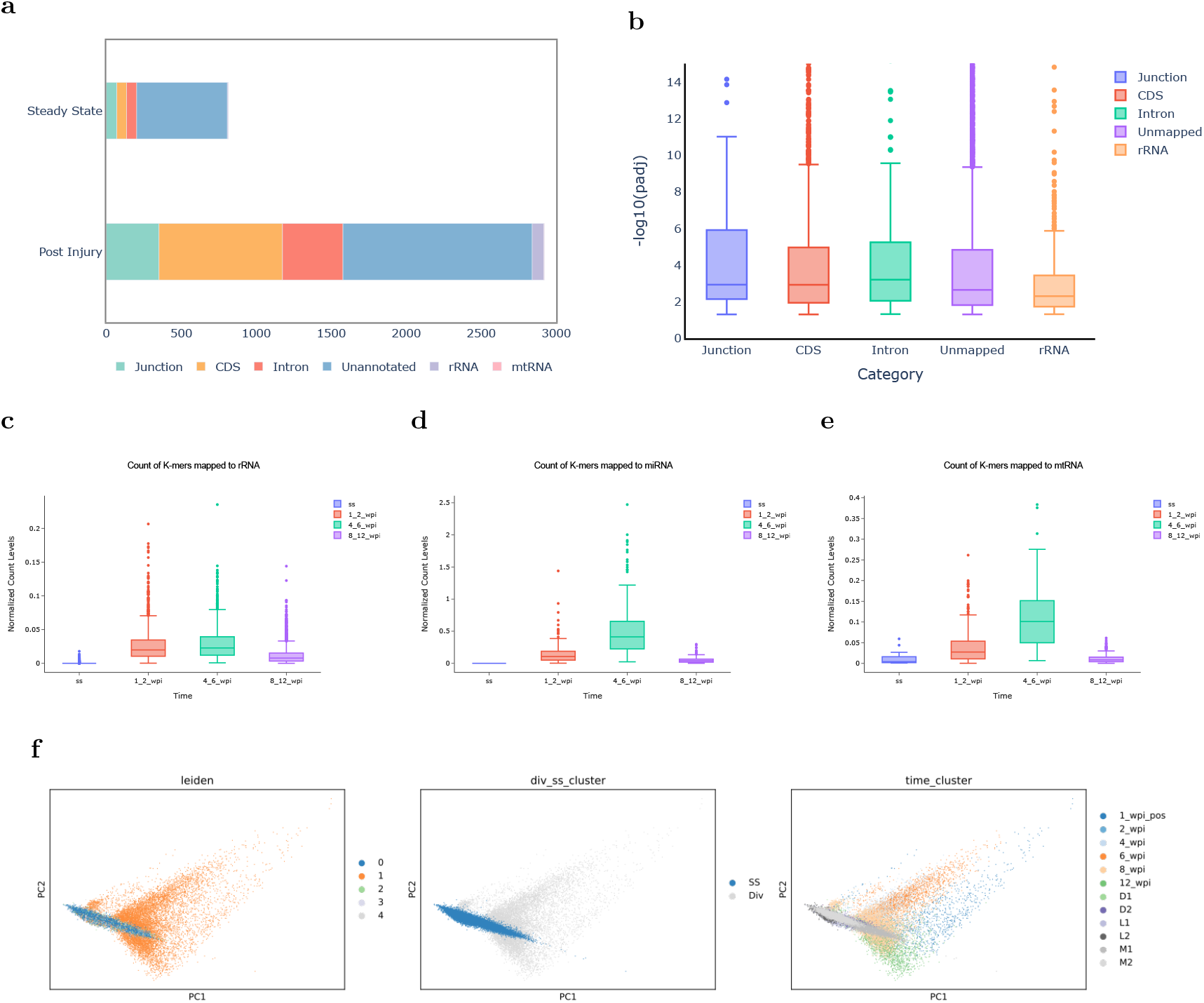
Axolotl Neuroregeneration Analysis: **a**. Numbers of contigs mapped to introns, junctions, CDS regions of the axolotl genome, as well as rRNA and mtRNA using homology search. A significant portion of contigs remained unannotated in both steady-state and post-injury conditions. Homology study was conducted on the unannotated contigs only. **b**. Distribution of p-values for *k*-mers within contigs from various annotations during post-injury phases. **c**. Average normalized counts of *k*-mers associated with rRNAs across time points, showing increased expression in post-injury time compared to steady-state. **d**. Elevated abundance of *k*-mers corresponding to miRNAs (*mir6236*) observed at weeks 4 and 6 post-injury. **e**. Increased abundance of *k*-mers related to mtRNA at weeks 1, 2, 4, and 6 post-injury, compared to steady-state and the later healing phase. **f**. PCA plots of identified clusters by utilizing the Leiden method on differentially expressed *k*-mer abundance matrix captured by scKAR highlighted according to **i**. Leiden clusters, **ii**. steady-state (ss) and post-injury (div) **iii**. timeline of post-injury (1 week post-injury - 1 wpi pos, 2 wpi, 4 wpi, 6 wpi, 8 wpi, 12 wpi) and steady-state samples (D1, D2, L1, L2, M1 and M2).

We extracted the constituent *k*-mers for the contigs that mapped to rRNAs, mtRNAs or miRNAs during both steady state and post-injury phase. By calculating their average counts from the *k*-mer abundance matrix over the weeks, we observed higher levels of the *k*-mers corresponding to rRNAs, miRNAs, and mtRNAs during the post-injury phase compared to both the steady-state condition and the period when healing was nearly complete as presented in Figures 3c, 3d, and 3e. We find that counts of *k*-mers mapping to rRNAs are high in both 1, 2 weeks and 4, 6 weeks after the injury whereas those of miRNA and mtRNAs are specifically high 4, 6 weeks post-injury.

We employed the Leiden clustering algorithm on the abundance matrix of differentially expressed *k*-mers. The PCA plot (Figure 3f) reveals a distinct pattern where all steady-state cells align along a nearly linear trajectory, with their expression primarily driven by the first principal component. In contrast, the div-seq cells exhibit a more dispersed distribution. Notably, one of the clusters was predominantly composed of cells from the post-injury phase, whereas the remaining clusters consisted primarily of steady-state cells, as shown in Supplementary Figure S1. Finally, We used scKAR to analyze the data from each week separately. For each week, we identified differentially expressed *k*-mers and contigs for various cell types like GLUT/GABA, NB, and Epen cells. The proportion of differentially expressed *k*-mers in each week is depicted in Supplementary Figure S2. This distribution is consistent with the proportion of different types of cells as mentioned in (21).

### scKAR indicates increased lncRNAs and intron retention in heart disease related genes in diseased cardiomyocytes

The heart, as the first organ to develop in the embryo, undergoes complex morphogenesis and defects in this process lead to congenital heart disease (CHD) (35; 36). Despite advancements in therapies, many patients with CHD face premature death due to heart failure and noncardiac causes. To understand this disease’s progression, single-nucleus RNA sequencing was performed (37) on 157,273 nuclei from control hearts and hearts from CHD patients, including those with hypoplastic left heart syndrome (HLHS) and tetralogy of Fallot (TOF), as well as dilated and hypertrophic cardiomyopathies (DCM and HCM) providing a comprehensive phenotyping of CHD. The dataset comprised 73,153 cardiomyocyte cells, where three distinct cell types were identified, labeled as CM1 (healthy cells), CM2 (mildly stressed cells), and CM3 (cells found exclusively in diseased conditions) (37). CM1 cells were predominantly found in donor samples, while CM3 cells were primarily observed in diseased samples.

Running scKAR, we identified contigs associated with CM1 and CM3. The resulting contigs were mapped to the human reference genome using BLAT (25). The outcomes of this analysis are summarized in Figure 4. We find a notable increase in non-transcriptomic events, such as intron retention and lncRNA transcription, in diseased cardiomyocytes (CM3) compared to control cells (CM1). To assess the significance of these differences, we constructed a contingency table of counts for uniquely mapped intronic and lncRNA contigs. A chi-square test with Yate’s correction was performed, revealing statistically significant enrichment of intron retention and lncRNA transcription in CM3, with *p*-values *<* 0.0006 and *<* 0.0001, respectively. Detailed proportions and counts for these events are shown in Figure 4a.

**Figure 4:**
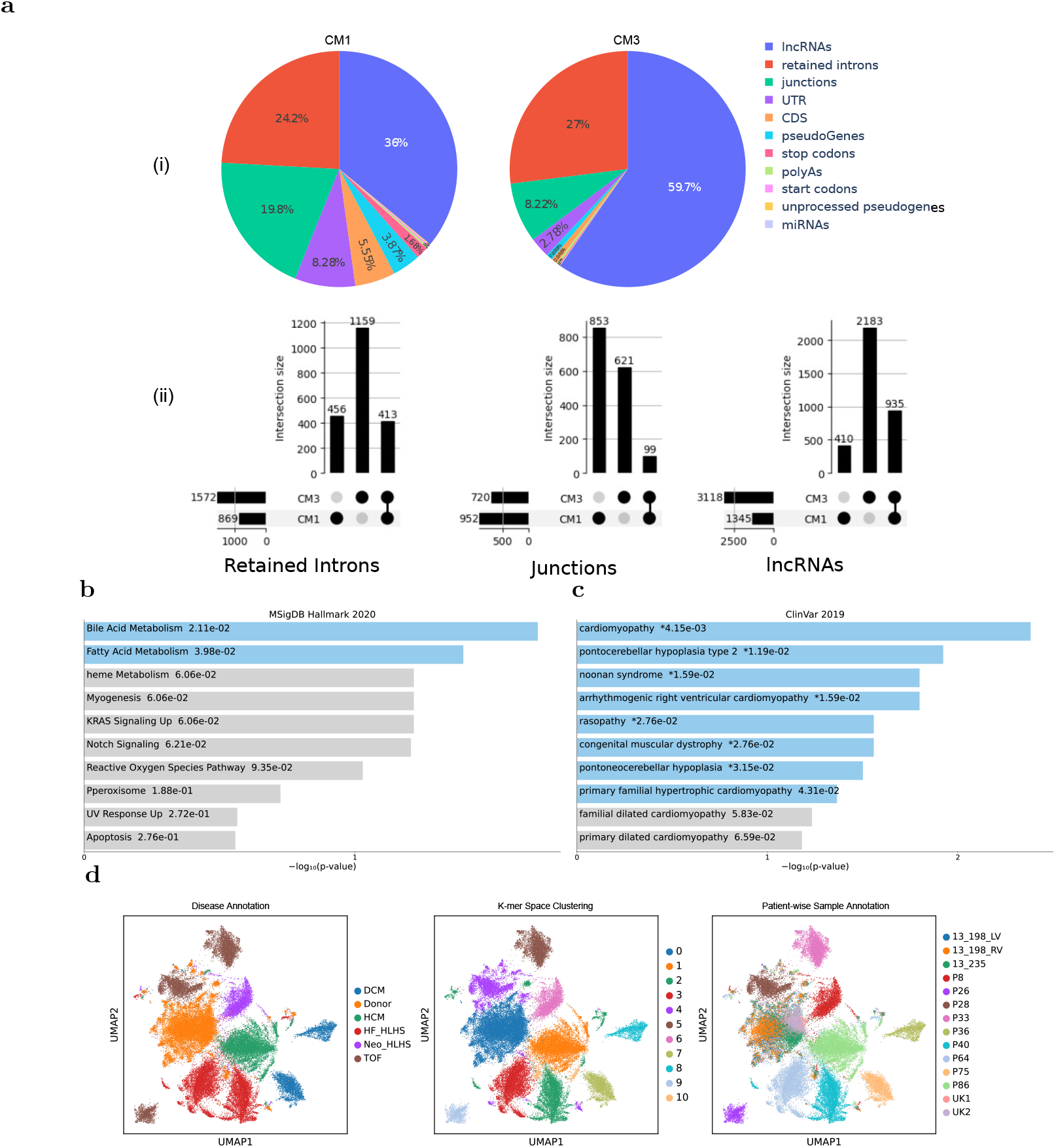
Analysis of congenital heart disease dataset: a.i. Breakdown of various transcriptional events in CM1 and CM3 samples. **a.ii**. Comparison of the numbers of genes corresponding to retained introns, junctions and lncRNAs in CM1, CM3 and both. **b** Enrichment analysis of genes associated with retained introns in CM1 samples using enrichR showing enrichment of genes in pathways related to metabolism, specifically bile acid and fatty acid metabolism. **c** Enrichment analysis of retained intron-associated genes in CM3 samples showing enrichment of genes in pathways implicated in cardiomyopathy, muscular dystrophy, and hypertrophic cardiomyopathy. **d** Leiden clustering on differentially expressed *k*-mer abundance matrix captured by scKAR, highlighted according to **i**. diagnosis type **ii**. Leiden clustering **iii**. sample ids: Case (P8, P26, P28, P33, P36, P40, P64, P75) and Control (13 198 LV, 13 198 RV, 13 235, UK1, UK2).

Given that splicing variants and intron retention are associated with various severe diseases (38; 39), we focused particularly on regions with retained introns, a relatively unexplored aspect of congenital heart disease. Stouffer’s method (40) was used to calculate the combined *p*-value for contigs derived from retained introns, based on the *p*-values of the constituent *k*-mers. By analyzing genes associated with low *p*-value contigs exhibiting intron retention, insights into the genes implicated in CM1 and CM3 were obtained. In CM1, intron retention likely stems from pre-spliced mRNA and occasional splicing errors, serving as a baseline. However, the elevated intron retention in CM3, despite lower read depth relative to CM1, suggests potential additional biological relevance.

For gene enrichment analysis, we ranked intron-mapping genes by increasing *p*-values and decreasing contig length, producing a prioritized gene set. The top 100 genes, shown in Supplementary Figures S6 and S5 for CM1 and CM3 respectively, were subjected to gene enrichment analysis using enrichR (41) using subsets of 20, 40, 60, 80, and 100 genes from the sorted list. The analysis for CM1 show enrichment of genes associated with metabolism and cardiovascular regulation such as TNNT2, HSF1, ACTN1, and LMNA, as shown in Figure 4b. In contrast, the gene list for CM3 reveal enrichment of genes linked to cardiomyopathy and cardiac regulatory processes, including TPM1, TNNI3K, CRIM1, PON2, and DSP, as shown in Figure 4c. The *p*-values and *q*-scores from enrichR are summarized in Supplementary Tables S2 and S3.

Additionally, lncRNAs and junction-contig mappings with low *p*-values were manually reviewed. Several identified lncRNAs mapped to OMIM genes associated with heart disease, underscoring the biological significance of these findings. Some of the notable instances can be found in Supplementary Table S7.

The final matrix of differentially expressed *k*-mer abundance was clustered to examine alignment with the metadata. Figure 4d illustrates that non-reference DE *k*-mers successfully distinguished between different cell and disease types. This suggests that reads traditionally unquantified by expression tools contain valuable diagnostic information from an embedding perspective.

## Discussion

This study introduces a robust, reference-free method, scKAR, for analyzing single-cell RNA sequencing (scRNA-seq) data, addressing critical limitations of conventional reference-based approaches. Our method, designed to circumvent the biases and constraints of incomplete or population-specific reference transcriptomes, excels in uncovering transcriptional events directly from raw data. By constructing *k*-mer abundance matrices and employing statistical models tailored for zero-inflated single-cell data, our approach provides an innovative solution for analyzing complex transcriptomic landscapes.

scKAR is particularly suited for studying non-model organisms, where reference transcriptomes are incomplete or entirely absent. We analyzed an axolotl neurogenesis dataset and identified elevated ribosomal RNA (rRNA) transcription during early regenerative phases and mitochondrial RNA (mtRNA) upregulation during peak neurogenesis. These findings align with the heightened metabolic and energy demands associated with tissue regeneration and neural development. Furthermore, our approach detected differential expression of *mir6236*, a microRNA implicated in regulating neural metabolism and development, emphasizing its sensitivity in identifying potentially biologically significant events undetected by reference-based approaches.

scKAR also demonstrates utility in detecting non-canonical transcriptional events—such as intron retention, long non-coding RNA (lncRNA) transcription, and atypical splicing—that are often overlooked by conventional pipelines. Moreover, the method is well-suited for examining stress or disease states, such as cancer, where rare transcriptional events play critical roles in disease progression. In an analysis of a congenital heart disease dataset, our tool successfully uncovered intron retention and lncRNA transcription of important genes for heart functions associated with cardiomyopathy. These findings underscore the method’s capacity to identify transcriptional anomalies in pathological conditions, highlighting potential avenues for further biological and clinical investigations.

Despite its strengths, the method has certain limitations. Statistical tests lose power in cases of data scarcity, making it challenging to identify subtle transcriptional events in sparsely sampled datasets. Additionally, clustering derived from the *k*-mer abundance matrix is computationally expensive, requiring substantial memory and potentially limiting its scalability for large datasets if gene expression matrix is unavailable. Future research could explore strategies to optimize partitioning techniques and reduce computational demands, thereby improving the method’s efficiency and applicability. Furthermore, extending the framework to detect gene-fusion events and quantify their relative abundance could provide additional insights into complex transcriptional processes.

## Methods

### Generating Normalized *k*-mer Abundance Matrix

The first step in scKAR is to generate a normalized *k*-mer abundance matrix. For this, the raw demultiplexed (cell-specific) sequence reads, provided in FASTA/FASTQ format, are processed using Jellyfish (42) to count 31-mers for each cell. *K*-mers with a count of 1 are discarded, as these are presumed to result from sequencing errors.

To account for variable read depth, transcript per million (TPM) normalization is applied to each cell. TPM normalization adjusts for differences in sequencing depth by considering the number of reads mapped per kilobase of transcript per million reads, allowing for comparisons of *k*-mer abundances both within and across samples. For *k*-mer abundance, TPM is computed using:

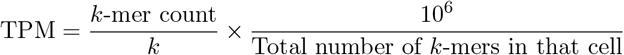

The normalized count vectors are subsequently merged to generate a cell-wise *k*-mer count matrix, referred to here as the *k*-mer abundance matrix. Due to the high level of zero inflation inherent in single-cell reads, only the non-zero values, along with their corresponding indices, were retained as an adjacency list for further processing.

### Filtering Reference *K*-mers

If a reference transcriptome is available, a reference *k*-mer list is precomputed from it. The *k*-mers present in the reference can be optionally filtered out from the *k*-mer abundance matrix for time and memory efficiency. The reference *k*-mers are stored in a sorted order to facilitate efficient filtering. To remove *k*-mers present in the reference list from the dataset, we employed a two-pointer filtering approach. This method operates in O(*n*) time complexity, where *n* represents the total number of unique *k*-mers identified across all cells.

### Partition Generation from Clustering

To generate partitions for identifying differentially expressed *k*-mers, we begin by clustering similar or homogeneous cell types using the gene expression matrix. The Leiden clustering algorithm (23), which is based on the Constant Potts Model (43) and uses a graph-based approach, is employed to detect clusters. The correlations among these clusters are used to construct a dendrogram, representing the hierarchical relationships among clusters as a binary tree, *T*, with root *R*. In this context, the leaves of the tree correspond to specific cell groups or conditions.

Let *U* be the set of all nodes and *E* be the set of all edges in the tree, *T*. Consider an edge, (*u, v*) *∈ E* whose removal splits the tree into two subtrees (trivially a node is also a subtree). Thur, the tree, *T* can be split into two groups: *X*, which is the set of all nodes belonging to the subtree connected to *u*, and the complementary group *U − X*. This produces a bipartition of the cell types or conditions, separating the most dissimilar clusters, as determined by their correlation in the dendrogram. Differential analysis is then performed on these partitions.

If the gene expression matrix is not available, clustering can be performed on the *k*-mer abundance matrix instead of the gene expression matrix. However, due to the large size of the *k*-mer abundance matrix, this requires significantly more physical memory. Users also have the option to provide custom bipartition-specific analyses based on predefined experiment conditions.

### Fisher’s F-test on *k*-mer Abundance Matrix

Given the large size of the *k*-mer abundance matrix, we focus on identifying *k*-mers that exhibit significant variance across different clusters or groups for narrowing down the *k*-mer space to test differential expression. Each cluster is treated as a separate group, and Fisher’s F-test (44) is applied to compare the variances between these groups.

**Null Hypothesis:** The means of all groups are equal.

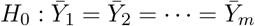

The F-statistic is computed as:

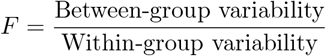

where the between-group variability is given by:

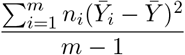

and the within-group variability is:

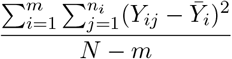

Here, *m* represents the number of groups, *n*_*i*_ is the sample size of the *i*-th group, 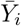 denotes the mean of the *i*-th group, and 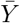 is the overall mean across all groups. *N* refers to the total number of observations, and *Y*_*ij*_ is the *j*-th observation in the *i*-th group.

The computed F-statistic is compared with the critical value from the F-distribution under the null hypothesis. To account for multiple comparisons, the Benjamini-Hochberg procedure is applied to control the false discovery rate (FDR) and adjust the p-values. We reject the null hypothesis for p-values ≤ 0.05, retaining only the *k*-mers with significant variance between groups.

### Detecting Differentially Expressed *k*-mers

The process of detecting differentially expressed *k*-mers involves four key stages: size factor estimation, linear model fitting, significance testing, and log-fold change computation. Singlecell expression matrices are highly zero-inflated, making traditional differential expression (DE) tools like DESeq2 (27), edgeR (46), and limma (47; 48), which assume a negative binomial distribution, unsuitable for single-cell assays. Adjustments to DESeq2 for single-cell data, as suggested by (28), have been experimentally validated in (49) (50).

For size factor estimation, we employ normalization by deconvolution (using the ‘Compute-Sum Factors’ function in R). The first step involves calculating the geometric mean for each *k*-mer:

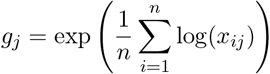

where *x*_*ij*_ is the *k*-mer count for cell *i* and *k*-mer *j*, and *n* is the total number of cells.

Next, the size factor for each cell is computed as:

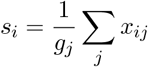

Normalization of the *k*-mer counts is then performed by adjusting the counts using the size factor:

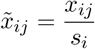

For model fitting, we utilize the generalized Gamma-Poisson model (50), which has been shown to better capture the sampling variability in single-cell data compared to negative binomial models (49). A lowered mean threshold appropriate for single-cell assays, is applied during the fitting process to filter out low-expression *k*-mers.

For statistical significance testing and log-fold change computation, the Likelihood Ratio Test (LRT) is used. The LRT compares two statistical models: the null model *H*_0_ and the alternative model *H*_1_, by evaluating the likelihoods *L*_0_ and *L*_1_ of the two models, respectively. The likelihood ratio is given by:

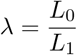

This is often expressed in terms of the log-likelihood ratio:

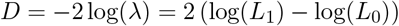

This statistic is used to assess the significance of differences in *k*-mer expression across conditions. Finally, The log fold change (logFC) is computed as the logarithm of the ratio of expression levels between the two conditions using:

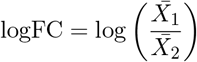

where 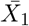 and 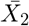 represent the mean expression values in the two conditions being compared. The two conditions are derived from the groups generated by internal edge splitting of the correlation dendrogram. Afterward, we employed the Benjamini-Hochberg correction (45) to control the false discovery rate (FDR). Finally, we identify differentially expressed *k*-mers by filtering for those with a logFC ≥ |1| and an FDR ≤ 0.05. For multiple internal edge splits, each bipartition of the dendrogram produces condition-specific *k*-mers that are differentially abundant in those specific conditions.

### Contig Generation

To generate contigs, the condition-specific *k*-mers are assembled using ABySS 2.0 (51) for de novo contig generation. The p-value for each contig is computed using Stouffer’s method for combining p-values (40) from the assembled constructs. For the 31-mers, contigs are considered valid if they exhibit at least a 27-mer overlap among the constructs.

### Post-Analysis

Several types of downstream analysis can be conducted based on the differentially expressed *k*-mers or contigs. Subpopulation within the samples can be uncovered by applying clustering techniques to the *k*-mer abundance matrix or the contig expression matrix. In the latter case, contig expression is inferred from the abundance of the constituent *k*-mers. Additionally, splicing events like intron retention, lncRNAs, and junctions can be detected by using tools like BLAT, which aligns contigs to the reference genome. Enrichment analysis on the genes associated with these events can reveal enriched pathways relevant to metabolism, cell signaling, disease progression, etc. For non-model organisms, the contigs may be mapped to a well-annotated genome of a closely related species to identify conserved functional elements.

### Datasets

The datasets used in this study are available through public repositories. Sequencing data (FASTQ files) and corresponding metadata for **Congenital Heart Disease** samples are accessible in the GEO Database (https://www.ncbi.nlm.nih.gov/geo/) under accession numbers GSM6165791 to GSM6165820. **Axolotl Neurogenesis** data are available in the European Nucleotide Archive (ENA) browser (https://www.ebi.ac.uk/ena/browser/) under accession numbers E-MTAB-11638, E-MTAB-11662, E-MTAB-11665, and E-MTAB-11666. **Metastatic Renal Cell Carcinoma** sequencing data can be accessed through the GEO Database under accession number GSE73121.

## Notes

### Competing Interest Statement

The authors have declared no competing interest.

