## Supplementary Information for "Reference-free Analysis of scRNA-seq Data Reveals Elevated rRNA and mtRNA Transcription during Neurogenesis in Axolotl"

| Contig ID | Accession ID | Species | Annotation | Bit Score | E-value |
|---|---|---|---|---|---|
| A_2099 | NC_006887 | *Ambystoma tigrinum* | Mitochondrion Complete Genome | 102.686 | 4.06705e-18 |
| A_3806 | MN135489 | *Ambystoma texanum* | Ribosomal RNA gene | 65.753 | 2.28719e-07 |
| A_877 | JQ511820 | *Rana pipiens* | Ribosomal RNA Gene | 76.8329 | 1.52623e-10 |
| A_650 | MW194271 | *Cynoglossus melampetalus* | Subunit Ribosomal RNA Gene | 78.6796 | 4.2435e-11 |
| A_1323 | MW624199 | *Cardioglossa nigromaculata* | Subunit Ribosomal RNA Gene | 58.3664 | 4.67778e-05 |
| A_3043 | XR_008814114 | *Lampris incognitus* | rRNA | 62.0597 | 2.79429e-06 |
| B_606 | XR_007812726 | *Lates calcarifer* | rRNA | 73.1396 | 1.7465e-09 |
| B_219 | XR_009323687 | *Acipenser ruthenus* | rRNA | 65.753 | 2.54132e-07 |
| B_589 | XR_010887693 | *Apteryx mantelli* | rRNA | 62.0597 | 2.95866e-06 |

Table S1: Blast Output of Unannotated Contigs during Post Injury and Steady State

| Enrichment Tag | p-value | q-value |
|---|---|---|
| Bile Acid Metabolism | 0.0211 | 0.1346 |
| Fatty Acid Metabolism | 0.0398 | 0.1346 |

Table S2: Enrichment analysis for CM1 condition genes (top 40 genes)

| Enrichment Tag | p-value | q-value |
|---|---|---|
| Cardiomyopathy | 0.0042 | 0.0397 |
| Pontocerebellar Hypoplasia Type 2 | 0.0119 | 0.0397 |
| Noonan Syndrome | 0.0159 | 0.0397 |
| Arrhythmogenic Right Ventricular Cardiomyopathy | 0.0159 | 0.0397 |
| Rasopathy | 0.0276 | 0.0451 |
| Congenital Muscular Dystrophy | 0.0276 | 0.0451 |
| Pontoneocerebellar Hypoplasia | 0.0315 | 0.0451 |
| Primary Familial Hypertrophic Cardiomyopathy | 0.0431 | 0.0539 |

Table S3: Enrichment analysis for CM3 condition genes (top 40 genes)

**a**

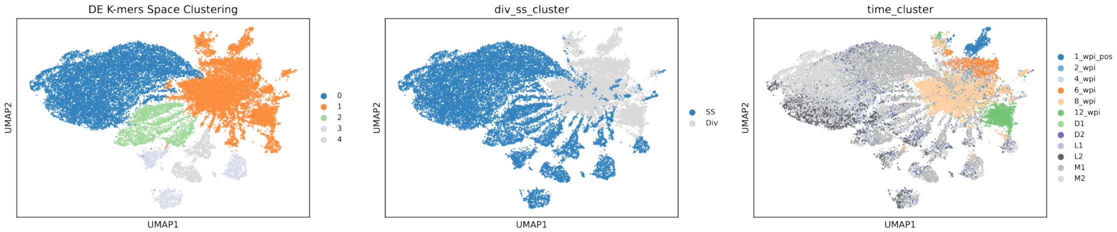


**b**

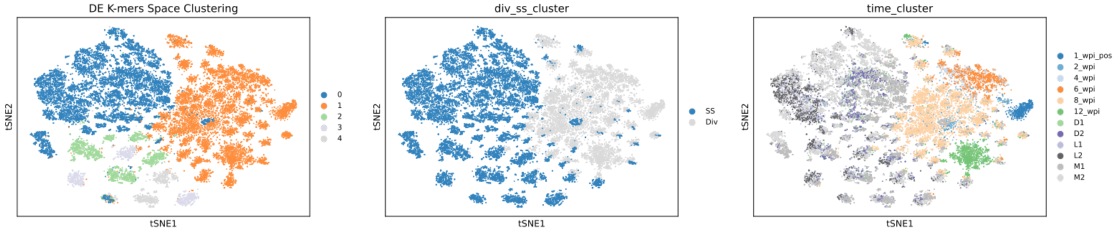


Figure S1: **a.** Axolotl Umap Clustering on Differentially Expressed K-mer Abundance Space. **b.** Axolotl T-SNE Clustering on Differentially Expressed K-mer Abundance Space.

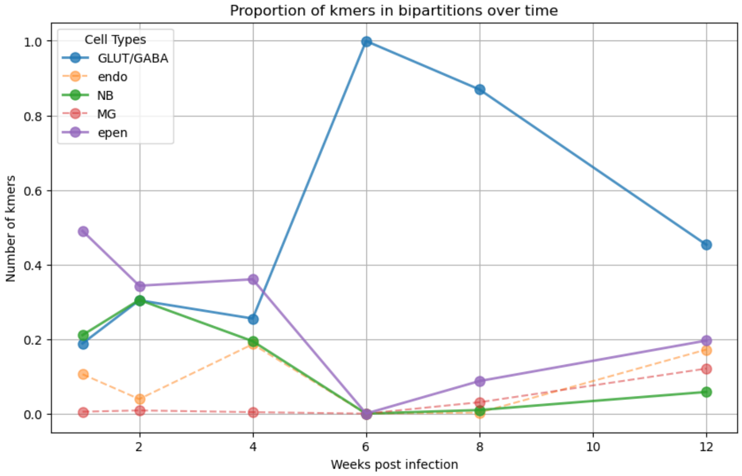


Figure S2: The normalized count of $k$-mers in different cell types in axolotl's neurogenesis phases.

| Contig ID | Gene Name | Disease |
|---|---|---|
| 3286 | TPM1 | Cardiomyopathy, DCM |
| 1965 | DMD | DCM |
| 1657 | DSP | DCM |
| 325 | CRIM1 | Marker for Diseased CMs |
| 1965 | MAP2K1 | Cardiac Hypertrophy |
| 859 | SH3GL3 | HCM |
| 3567 | TNNI3K | HCM |

Table S4: Genes Associcated with Intron Retention in CM3 samples

| Contig ID | Chromosome | Start | End | Gene Name |
|---|---|---|---|---|
| A_1965 | chr12 | 44553022 | 44553069 | NELL2 |
| A_1946 | chr3 | 197625669 | 197625758 | SDHAP4 |
| A_1965 | chr15 | 66452040 | 66452087 | MAP2K1 |
| A_3286 | chr15 | 63066076 | 63066018 | TPM1 |
| A_1965 | chr5 | 31203507 | 31203460 | CDH6 |
| A_1965 | chr12 | 55690872 | 55690919 | ITGA7 |
| A_1965 | chrX | 31530935 | 31530888 | DMD |
| A_1657 | chr6 | 7554882 | 7554840 | DSP |
| A_1965 | chr7 | 30445014 | 30445061 | NOD1 |
| A_859 | chr15 | 83518737 | 83518716 | SH3GL3 |
| A_1965 | chr1 | 74445480 | 74445433 | TNNI3K |
| A_1965 | chr18 | 63363481 | 63363434 | KDSR |
| A_1965 | chr20 | 20314796 | 20314749 | CFAP61 |
| A_2318 | chr8 | 105896352 | 105896388 | ZFPM2-AS1 |
| A_1965 | chr12 | 21287764 | 21287811 | SLCO1A2 |
| A_1423 | chr7 | 95425191 | 95425134 | PON2 |
| A_3567 | chr22 | 28669120 | 28669154 | TTC28 |
| A_1965 | chr15 | 64332427 | 64332474 | ENSG00000259316 |
| A_1965 | chr15 | 60954195 | 60954242 | RORA |
| A_1965 | chr10 | 31443672 | 31443625 | ZEB1 |
| A_1965 | chr2 | 71411735 | 71411688 | ZNF638 |
| A_1965 | chr9 | 41155736 | 41155783 | ZNG1F |
| A_1965 | chr11 | 35395936 | 35395889 | SLC1A2 |
| A_1965 | chrX | 11219147 | 11219100 | ARHGAP6 |
| A_1965 | chr7 | 14709719 | 14709766 | DGKB |
| A_1942 | chr9 | 665185 | 665156 | KANK1 |
| A_1941 | chr7 | 131097411 | 131097559 | LINC-PINT |
| A_1938 | chrX | 9079022 | 9078993 | FAM9B |
| A_1938 | chr2 | 85640743 | 85640778 | USP39 |
| A_1938 | chr17 | 69248987 | 69249025 | ABCA5 |
| A_1938 | chr4 | 40945986 | 40945949 | APBB2 |
| A_1938 | chr17 | 39142895 | 39142852 | PLXDC1 |
| A_1919 | chr1 | 151809179 | 151809128 | RORC |
| A_1919 | chr3 | 134541019 | 134541069 | CEP63 |
| A_1946 | chr3 | 195988079 | 195988168 | SDHAP1 |
| A_1919 | chr21 | 46350005 | 46349907 | PCNT |
| A_1965 | chr15 | 83451065 | 83451018 | SH3GL3 |
| A_1965 | chr5 | 14809088 | 14809135 | ANKH |
| | | | | *Continued on next page* |

| Contig ID | Chromosome | Start | End | Gene Name |
|---|---|---|---|---|
| A_1965 | chr17 | 594062 | 594109 | VPS53 |
| A_1965 | chr22 | 28669055 | 28669102 | TTC28 |
| A_1965 | chr13 | 59848576 | 59848529 | DIAPH3 |
| A_1965 | chr12 | 9582193 | 9582240 | ENSG00000284634 |
| A_1965 | chr8 | 47332259 | 47332212 | SPIDR |
| A_1965 | chr7 | 70203097 | 70203144 | AUTS2 |
| A_1965 | chr13 | 77204801 | 77204848 | MYCBP2 |
| A_1965 | chr6 | 52478178 | 52478225 | EFHC1 |
| A_1965 | chr13 | 52059707 | 52059754 | NEK5 |
| A_1965 | chr7 | 33548607 | 33548560 | BBS9 |
| A_1965 | chr7 | 37298563 | 37298516 | ELMO1 |
| A_1965 | chr17 | 30439668 | 30439621 | CPD |
| A_1965 | chr17 | 32678518 | 32678471 | MYO1D |
| A_1965 | chr17 | 56193155 | 56193202 | ANKFN1 |
| A_1965 | chr20 | 58687229 | 58687182 | STX16-NPEPL1 |
| A_1965 | chr5 | 5216866 | 5216819 | ADAMTS16 |
| A_1965 | chr7 | 18135562 | 18135515 | HDAC9 |
| A_1897 | chr12 | 29725606 | 29725502 | TMTC1 |
| A_1892 | chr17 | 32050195 | 32050235 | LRRC37B |
| A_1872 | chr11 | 114184872 | 114184766 | ZBTB16 |
| A_1657 | chr19 | 38215785 | 38215760 | DPF1 |
| A_1657 | chr16 | 31144543 | 31144506 | PRSS36 |
| A_1657 | chr4 | 26348938 | 26348905 | RBPJ |
| A_1657 | chr12 | 82869623 | 82869656 | TMTC2 |
| A_1657 | chr1 | 39355922 | 39355876 | MACF1 |
| A_1657 | chr17 | 65758069 | 65758032 | CEP112 |
| A_1657 | chr10 | 66940441 | 66940480 | CTNNA3 |
| A_1657 | chr8 | 100010879 | 100010834 | RGS22 |
| A_1657 | chr19 | 13450894 | 13450856 | CACNA1A |
| A_1657 | chr7 | 74889576 | 74889522 | ENSG00000290833 |
| A_817 | chr21 | 46351540 | 46351427 | PCNT |
| A_843 | chr1 | 11845887 | 11845943 | CLCN6 |
| A_1657 | chr7 | 75958762 | 75958806 | POR |
| A_1657 | chr5 | 95513428 | 95513384 | SKIC3 |
| A_1657 | chr1 | 42544128 | 42544076 | CCDC30 |
| A_856 | chr10 | 175285 | 175241 | ZMYND11 |
| A_859 | chr5 | 31925492 | 31925643 | PDZD2 |
| A_859 | chr1 | 169300965 | 169301022 | NME7 |
| A_859 | chr6 | 101600034 | 101599970 | GRIK2 |
| A_859 | chr14 | 67841455 | 67841401 | RAD51B |
| A_859 | chr7 | 92182797 | 92182748 | ENSG00000289027 |
| A_1657 | chr11 | 46470224 | 46470268 | AMBRA1 |
| A_1657 | chr7 | 70395269 | 70395289 | AUTS2 |
| A_1697 | chr5 | 68291178 | 68291062 | PIK3R1 |
| A_1707 | chr11 | 114184784 | 114184702 | ZBTB16 |
| A_3567 | chr17 | 82775709 | 82775772 | TBCD |
| A_1826 | chr12 | 29745396 | 29745261 | TMTC1 |
| A_1795 | chr16 | 6703412 | 6703423 | RBFOX1 |
| A_1795 | chr17 | 38374388 | 38374399 | SOCS7 |
| A_1795 | chr12 | 6332283 | 6332298 | TNFRSF1A |

*Continued on next page*

| Contig ID | Chromosome | Start | End | Gene Name |
|---|---|---|---|---|
| A_1795 | chr14 | 58498354 | 58498331 | KIAA0586 |
| A_1795 | chr8 | 129966871 | 129966845 | CYRIB |
| A_1795 | chr18 | 23321635 | 23321601 | TMEM241 |
| A_1795 | chr6 | 161931283 | 161931305 | PRKN |
| A_1795 | chr19 | 48902579 | 48902553 | NUCB1 |
| A_1795 | chr19 | 13370638 | 13370607 | CACNA1A |
| A_1795 | chr9 | 109025349 | 109025313 | TMEM245 |
| A_1795 | chr16 | 21716311 | 21716350 | OTOA |
| A_1795 | chr1 | 67113590 | 67113550 | C1orf141 |
| A_1795 | chr5 | 14266312 | 14266269 | TRIO |
| A_1795 | chr15 | 90605054 | 90605102 | CRTC3 |
| A_1795 | chr8 | 33487677 | 33487626 | MAK16 |

Table S5: Genes associated with upregulated contigs mapping to retained introns in CM3

| Contig ID | Chromosome | Start | End | Gene Name |
|---|---|---|---|---|
| B_0 | chr4 | 94466472 | 94466102 | PDLIM5 |
| B_1202 | chr7 | 154457854 | 154457910 | DPP6 |
| B_1201 | chr8 | 41504002 | 41504058 | GOLGA7 |
| B_1200 | chr8 | 47332130 | 47332074 | SPIDR |
| B_1199 | chr2 | 36874486 | 36874543 | STRN |
| B_1198 | chr6 | 85601864 | 85601807 | ENSG00000271793 |
| B_1197 | chr7 | 80674275 | 80674218 | CD36 |
| B_1196 | chr8 | 144314451 | 144314396 | HSF1 |
| B_1195 | chr1 | 11998669 | 11998613 | MFN2 |
| B_1194 | chr14 | 101877194 | 101877138 | PPP2R5C |
| B_1193 | chr20 | 35712232 | 35712288 | RBM39 |
| B_1192 | chr17 | 7014385 | 7014330 | RNASEK |
| B_1191 | chr17 | 7014385 | 7014330 | RNASEK-C17orf49 |
| B_1190 | chr13 | 32523451 | 32523507 | N4BP2L2 |
| B_1189 | chr6 | 128904710 | 128904766 | LAMA2 |
| B_1188 | chrX | 131795516 | 131795574 | FIRRE |
| B_1187 | chr16 | 7375783 | 7375726 | RBFOX1 |
| B_1186 | chr1 | 26454691 | 26454750 | DHDDS |
| B_1185 | chr4 | 185718003 | 185718059 | SORBS2 |
| B_1184 | chr19 | 47779587 | 47779531 | SELENOW |
| B_1183 | chr7 | 75146248 | 75146191 | GTF2IRD2B |
| B_1182 | chr12 | 48129453 | 48129510 | PFKM |
| B_1181 | chr4 | 83593370 | 83593313 | GPAT3 |
| B_1180 | chr12 | 20537694 | 20537636 | PDE3A |
| B_1203 | chr8 | 99220919 | 99220863 | VPS13B |
| B_1204 | chr7 | 129867581 | 129867637 | UBE2H |
| B_1205 | chr7 | 111969092 | 111969148 | DOCK4 |
| B_1206 | chr7 | 70203226 | 70203282 | AUTS2 |
| B_1230 | chr14 | 95138849 | 95138905 | DICER1 |
| B_1229 | chr3 | 183234828 | 183234772 | MCF2L2 |
| B_1228 | chr15 | 51583257 | 51583201 | DMXL2 |
| B_1227 | chr6 | 65550469 | 65550413 | EYS |
| B_1226 | chr6 | 125289363 | 125289419 | HDDC2 |
| B_1225 | chr22 | 28669184 | 28669240 | TTC28 |
| B_1224 | chr6 | 129004285 | 129004341 | LAMA2 |
| B_1223 | chr1 | 35484936 | 35484880 | KIAA0319L |
| B_1222 | chr1 | 41208308 | 41208364 | SCMH1 |
| B_1221 | chr4 | 78111634 | 78111690 | FRAS1 |
| B_1220 | chr1 | 74448142 | 74448198 | TNNI3K |
| B_1179 | chr13 | 45337324 | 45337381 | TPT1 |
| B_1219 | chr18 | 62096662 | 62096606 | PIGN |
| B_1217 | chr4 | 7994229 | 7994173 | ABLIM2 |
| B_1216 | chr20 | 23432009 | 23432065 | CSTL1 |
| B_1215 | chr2 | 196911488 | 196911544 | PGAP1 |
| B_1214 | chr5 | 119605333 | 119605389 | HSD17B4 |
| B_1213 | chr2 | 115488762 | 115488818 | DPP10 |
| B_1212 | chr2 | 61979879 | 61979823 | COMMD1 |
| B_1211 | chr2 | 112503944 | 112503888 | TTL |
| B_1210 | chr11 | 99607982 | 99608038 | CNTN5 |

*Continued on next page*

| Contig ID | Chromosome | Start | End | Gene Name |
|---|---|---|---|---|
| B_1209 | chr11 | 85500423 | 85500479 | DLG2 |
| B_1208 | chr8 | 113060124 | 113060068 | CSMD3 |
| B_1207 | chr7 | 30445143 | 30445199 | NOD1 |
| B_1218 | chr4 | 87347255 | 87347199 | HSD17B11 |
| B_1178 | chr19 | 1272958 | 1272902 | CIRBP |
| B_1177 | chr11 | 46992837 | 46992894 | CSTPP1 |
| B_1176 | chr11 | 134089908 | 134089965 | JAM3 |
| B_1146 | chr15 | 83453900 | 83453959 | SH3GL3 |
| B_1145 | chr11 | 85327694 | 85327635 | DLG2 |
| B_1144 | chr11 | 99605020 | 99604961 | CNTN5 |
| B_1143 | chr11 | 87050237 | 87050178 | TMEM135 |
| B_1142 | chr14 | 30687906 | 30687965 | SCFD1 |
| B_1141 | chr7 | 30442178 | 30442119 | NOD1 |
| B_1140 | chr7 | 70200263 | 70200204 | AUTS2 |
| B_1139 | chr5 | 79782009 | 79782068 | CMYA5 |
| B_1138 | chr5 | 9259813 | 9259754 | SEMA5A |
| B_1137 | chr6 | 129001320 | 129001261 | LAMA2 |
| B_1136 | chrX | 66183794 | 66183853 | HEPH |
| B_1147 | chr3 | 183237793 | 183237852 | MCF2L2 |
| B_1135 | chr6 | 133472186 | 133472245 | EYA4 |
| B_1133 | chr4 | 98595528 | 98595587 | TSPAN5 |
| B_1132 | chr4 | 78108670 | 78108611 | FRAS1 |
| B_1131 | chr3 | 98743267 | 98743209 | ST3GAL6 |
| B_1130 | chr4 | 87350212 | 87350271 | HSD17B11 |
| B_1129 | chr16 | 21268989 | 21269046 | CRYM |
| B_1128 | chr3 | 65949913 | 65949856 | MAGI1 |
| B_1127 | chr13 | 77201964 | 77201905 | MYCBP2 |
| B_1126 | chr7 | 14662133 | 14662192 | DGKB |
| B_1125 | chr9 | 71205141 | 71205200 | TRPM3 |
| B_1124 | chr17 | 74203735 | 74203678 | RPL38 |
| B_1123 | chr1 | 1634520 | 1634464 | MMP23B |
| B_1134 | chrX | 31533556 | 31533497 | DMD |
| B_1231 | chr1 | 201364316 | 201364371 | TNNT2 |
| B_1148 | chr5 | 81619024 | 81618965 | SSBP2 |
| B_1150 | chr19 | 29462225 | 29462166 | VSTM2B-DT |
| B_1175 | chr20 | 8533463 | 8533520 | PLCB1 |
| B_1174 | chr8 | 39200221 | 39200278 | ADAM32 |
| B_1173 | chr4 | 185766049 | 185766106 | SORBS2 |
| B_1172 | chr4 | 94524286 | 94524230 | PDLIM5 |
| B_1171 | chr4 | 163717596 | 163717653 | MARCHF1 |
| B_1170 | chr5 | 178024406 | 178024463 | FAM153CP |
| B_1169 | chr3 | 113479575 | 113479632 | SPICE1 |
| B_1168 | chr7 | 100315018 | 100315076 | SPDYE3 |
| B_1167 | chr17 | 59180919 | 59180861 | PRR11 |
| B_1165 | chr17 | 7014308 | 7014252 | RNASEK |
| B_1164 | chr17 | 7014308 | 7014252 | RNASEK-C17orf49 |
| B_1149 | chr2 | 196908523 | 196908464 | PGAP1 |
| B_1163 | chr7 | 134933448 | 134933391 | CALD1 |
| B_1161 | chr10 | 32305686 | 32305744 | EPC1 |
| B_1160 | chr10 | 17616440 | 17616497 | HACD1 |

*Continued on next page*

| Contig ID | Chromosome | Start | End | Gene Name |
|---|---|---|---|---|
| B_1159 | chr2 | 33530692 | 33530749 | RASGRP3 |

Table S6: Genes associated with upregulated contigs mapping to retained introns in CM1

| Contig ID | Chr | Start | End | Gene Name | Omim ID | Comments |
|---|---|---|---|---|---|---|
| A_4350 | chr1 | 237805747 | 237805904 | RYR2 | 180902 | Ventricular arrhythmias due to cardiac ryanodine receptor calcium release deficiency syndrome |
| A_102 | chr2 | 86214158 | 86214232 | REEP1 | 609139 | Neuronopathy, distal hereditary motor, autosomal dominant |
| A_23 | chr11 | 114254282 | 114254160 | ZBTB16 | 176797 | Leukemia, acute promyelocytic |
| A_258 | chr7 | 80624461 | 80624562 | CD36 | 173510 | Coronary heart disease |
| A_3273 | chr5 | 128723811 | 128723874 | ENSG00000248634 | 614106 | Myopathy with myalgia, increased serum creatine kinase, and with or without episodic rhabdomyolysis |

Table S7: Identified OMIM alleles by manually annotating contigs sorted by p-value
